# Cultivating microbial communities from the serpentinite-hosted Prony Bay Hydrothermal Field on different carbon sources in hydrogen-fed bioreactors

**DOI:** 10.1101/2025.06.11.659039

**Authors:** Rabja Maria Popall, Agathe Roland, Sylvain Davidson, Yannick Combet-Blanc, Roy E. Price, Marianne Quéméneur, Anne Postec, Gaël Erauso

**Affiliations:** Aix-Marseille Univ, Univ Toulon, CNRS, IRD, MIO UM 110, Marseille, France; Stony Brook University, SoMAS Stony Brook, New York 11794, USA

**Keywords:** serpentinization, submarine alkaline vent, hydrothermal system, alkaliphile, hydrogenotroph, primary production, carbon fixation, origin of life

## Abstract

The primary source of carbon is one of the most fundamental questions regarding the development of microbial communities in serpentinite-hosted systems. The hydration of ultramafic rock to serpentinites generates hydrogen and creates hyper-alkaline conditions that deplete the environment of dissolved inorganic carbon. Metagenomic studies suggest that serpentinite-hosted microbial communities depend on the local redissolution of bicarbonate as well as on small organic molecules produced by abiotic reactions associated with serpentinization. To test these hypotheses, microbial consortia collected from the Prony Bay hydrothermal field were grown under anoxic conditions in hydrogen-fed bioreactors using bicarbonate, formate, acetate, or glycine as the sole carbon source. In contrast to glycine, the other three carbon substrates allowed the growth of microbial consortia characterized by significant enrichment of individual taxa. Surprisingly, these taxa were dominated by microbial genera characterized as aerobic rather than anaerobic as expected. We propose that an intricate feedback loop between autotrophic and heterotrophic foundation species facilitates the establishment of serpentinite-hosted shallow subsurface ecosystems. Bicarbonate-fixing *Meiothermus* and *Hydrogenophaga*, as well as formate-fixing *Meiothermus*, *Thioalkalimicrobium,* and possibly a novel genotype of *Roseibaca* might produce organic compounds for heterotrophs at the first trophic level. In addition, the base of the trophic network may include heterotrophic *Roseibaca*, *Acetoanaerobium,* and *Meiothermus* species producing CO_2_ from acetate for a more diverse community of autotrophs. The cultivated archaeal community is expected to recycle CH_4_ and CO_2_ between *Methanomicrobiales* and *Methanosarcinales* with putative *Woesearchaeales* symbionts.

## Background

Hydrothermal vents driven by serpentinization are a window into the past, possibly all the way back to the origin of life. Serpentinization, the hydration of exposed mantle rock, creates geochemical conditions that most likely characterized early Earth [1]. Most modern organisms cannot survive in these conditions, except for specialized microbial communities that grow independently of oxygen (O_2_). Elucidating their metabolism may be key to the first life forms and the current limits of life [2].

The hydration of mantle rocks to serpentinites produces hydrothermal fluids enriched in hydroxide ions and hydrogen (H_2_). While the released H_2_ constitutes a potent energy source for microbial metabolism, the elevated pH poses a fundamental energetic challenge to life. Firstly, it inverts the cell membrane potential, requiring particular adaptations to maintain the energetic driving force of the cell [3]. It has been shown that some alkaliphiles use Na^+^ instead of proton gradients or can concentrate protons at the cell surface [4]. Secondly, the elevated pH reduces the solubility of essential nutrients such as phosphorous [5] and dissolved inorganic carbon (DIC). The latter is usually vital for primary production, which constitutes the base of the trophic network. In hyperalkaline environments, however, carbon dioxide (CO_2_) precipitates as calcium carbonate (CaCO_3_), forming carbonaceous hydrothermal chimneys over time and removing DIC from the pool of bioavailable carbon [6]. This phenomenon raises one of the main questions regarding the functioning of serpentinite-hosted microbial ecosystems: which carbon source can replace CO_2_ in primary production?

Various small organic acids and amino acids produced abiotically by serpentinization may hold the key to this issue. Formate and acetate are yielded in Fischer-Tropsch and Sabatier-type reactions catalyzed in the hydrothermal fluid [3, 7, 8]. Similarly, serpentinizing conditions have been shown to facilitate the abiotic synthesis of glycine [9, 10]. Formate, acetate, and glycine concentrations vary widely between hydrothermal systems [7, 11], but all three compounds are likely ubiquitous in serpentinite-hosted environments. In addition, some serpentinite-hosted microorganisms have been shown to grow on solid CaCO_3_. The physiological mechanism they employ probably involves the local redissolution of bicarbonate ions [4]. For organisms capable of such redissolution, the carbonaceous hydrothermal chimneys could provide a vast carbon source.

By definition, primary producers are autotrophic and use an inorganic carbon source that is converted into more complex carbohydrates via a carbon fixation pathway. For this reason, acetate and glycine in particular, but also formate, are not usually considered as primary carbon sources. Due to the lack of DIC, however, the concept of primary production in serpentinite-hosted ecosystems might need to be reconsidered. This is supported by the fact that all of the compounds mentioned above are derived from an abiotic source, which *per se* challenges the definitions of heterotrophy and autotrophy [12].

In this context, formate and bicarbonate can be interpreted as autotrophic carbon sources. Once accumulated in the pH-neutral cytoplasm, they are both converted into CO_2_ by the formate dehydrogenase or the carbonic anhydrase, respectively [13, 14]. The yielded CO_2_ can then be used in any of the seven known carbon fixation pathways to form acetyl-CoA or other precursors for respiration, fermentation, and heterotrophic metabolism. On the contrary, acetate and glycine cannot be metabolized through a canonical carbon fixation pathway and thus constitute heterotrophic carbon sources. Acetate is directly converted to acetyl-CoA by the acetyl-CoA synthetase or the acetate kinase and phosphate acetyltransferase [15, 16], while glycine is reduced to acetyl phosphate by the glycine reductase complex. The only known exception to this heterotrophic route is syntrophic acetate oxidation (SAO), where a symbiotic bacterium oxidizes acetate to CO_2_ and H_2_ or formate for an autotrophic methanogen or sulfate-reducer. However, SAO has only been described in a handful of taxa [17]. While heterotrophs are not typically regarded as primary producers, they might play a more fundamental role in the serpentinite-hosted trophic network than elsewhere. Lang et al. [18] proposed that heterotrophic foundation species could produce CO_2_ for a diverse community of autotrophs, including those without formate or bicarbonate uptake genes. Acetate- and glycine assimilating heterotrophs could thus be as essential for the trophic chain as bicarbonate- or formate-fixing autotrophs.

In the last decade, the metabolic potential of these hypothetical primary producers has mainly been explored through metagenomic approaches, leading to some exciting discoveries of novel routes for non-canonical carbon fixation pathways and hypotheses on ecosystem functioning [4, 10, 14, 15]. However, many of the concerned enzymes are bidirectional or have other metabolic functions, rendering it difficult to evaluate such hypotheses based on metagenomic predictions alone. This requires experimental evidence, for example by observing the metabolic activity of targeted communities in culture. However, the cultivation of serpentinite-hosted communities has proven problematic, mainly because the physicochemical conditions associated with serpentinization are challenging to reproduce in the laboratory. Previous attempts have focused on microcosm experiments, but produced contradictory results: Kohl et al. [19] observed methane (CH_4_) production from bicarbonate, formate and acetate in microbial communities from The Cedars ophiolite, while the same experiment failed with microbial communities from the Tablelands ophiolite [20]. To bridge this gap, we performed cultivation experiments on natural serpentinite-hosted communities in controlled laboratory conditions. These communities were recovered from the shallow Prony Bay hydrothermal field [21] and enriched on a bioreactor platform, which allowed the control of various environmental parameters such as pH, temperature, and continuous gas supply (Figure 1). Four hydrogenotrophic culture conditions were established, each alimented with bicarbonate, formate, acetate, or glycine as the sole carbon source. Carbon consumption, growth, diversity, and taxonomic composition were compared between the microbial consortia to determine which carbon sources facilitate the development of microbial communities in serpentinizing conditions and which taxonomic groups are enriched on them.

**Figure 1.**
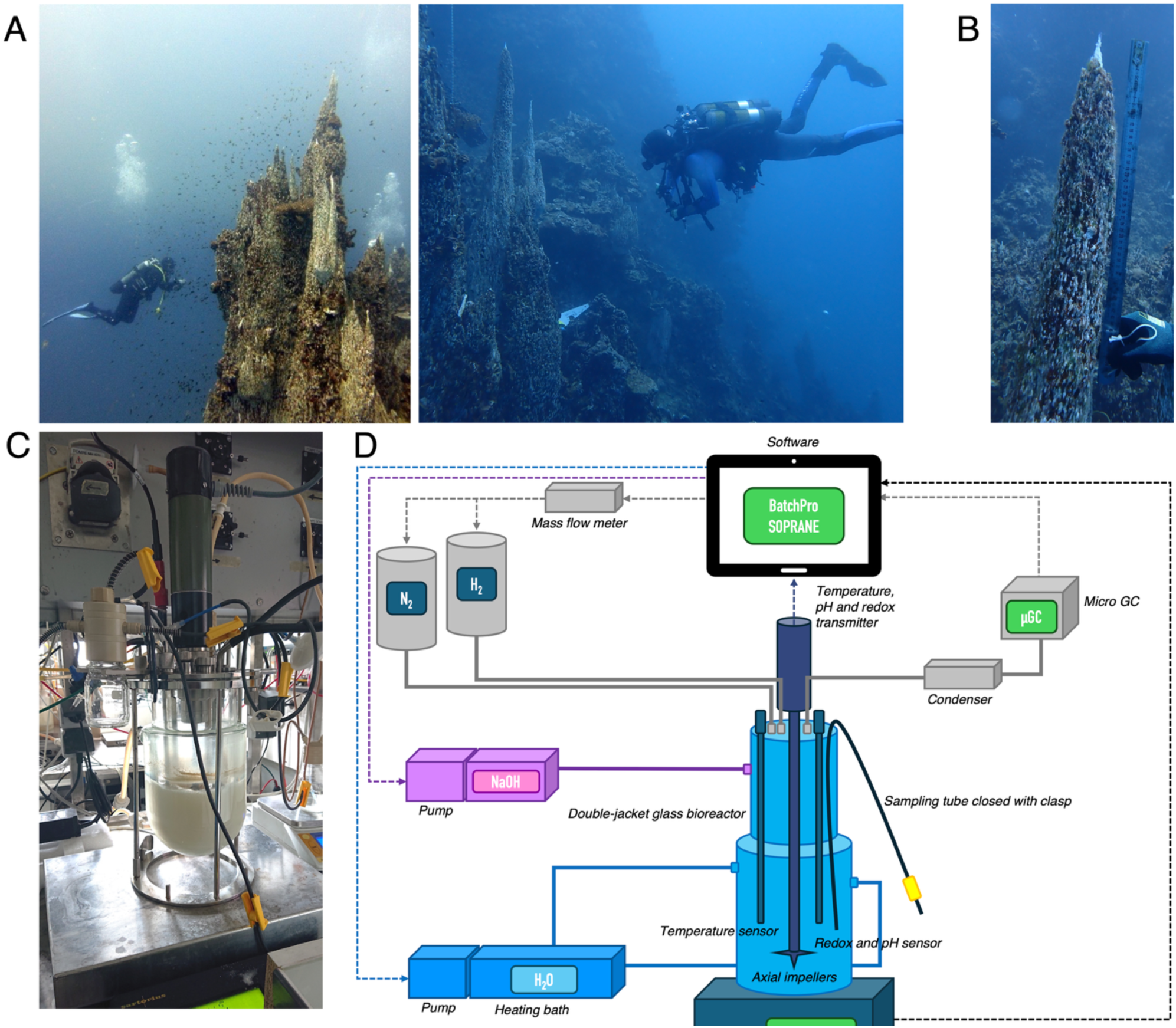
A) The “Aiguille de Prony” which measures 38 m from top to bottom. The exact study site is shown in the photo to the right. B) The chimney collected from the “Aiguille de Prony”. C) Bioreactor used in cultivation experiments with a working volume of 1.5 L. D) Schematic illustration of the bioreactor set-up.

## Material and methods

### Study site

The shallow Prony Bay hydrothermal field is located at the southern end of New Caledonia. As a coastal system, it comprises several sites from land to sea that reflect the geomicrobiological characteristics of terrestrial and marine serpentinite-hosted systems, respectively [5, 22–25]. The most prominent hydrothermal mount is the “Aiguille de Prony”, a large carbonate massif (38 m high from its base at a depth of around 45 m, depending on the tide, to its summit) (Figure 1A). The most active zone lies at a depth of 20 meters, where numerous chimneys emit hydrothermal fluid. The concentration of DIC measured in the fluid remains below 1 mmol L^-1^, with pH values ranging from 10 to 11.3, Eh values ranging from - 540 to -680 mV, and a temperature of around 35°C measured in the endmember fluid [21, 26]. During a field campaign in 2022, the submittal part (∼ 50 cm long and max 20 cm in diameter) of a mature, active chimney was sampled. The chimney was covered with a watertight bag, letting the hydrothermal fluid accumulate for a few hours until the ambient seawater was entirely removed. The chimney was then sawed off, the bag hermetically sealed and transported to the field laboratory. The chimney was divided into slices of approximately 35 mm under sterile conditions. The outer layer was removed with a chisel to isolate the center of each slice. This part was ground with mortar and pestle in an anaerobic chamber. The resulting slurry was aliquoted into hermetically sealed Schott bottles and stored under a nitrogen (N_2_) atmosphere at 4-8°C until inoculation of cultures.

### Experimental set-up

Experiments were carried out in sterile, non-opaque double-jacketed glass bioreactors (Pierre Guerin, France) with a working volume of 1.5 L (Figure 1B). The cultures were agitated at 500 rpm, the temperature was continuously montitored by a probe (Prosensor pt 100, France) and maintained at 35 ± 2 °C via hot water circulation in the double jacket via a thermostat (Julabo, SE6, Switzerland). The bioreactors were supplied with a continuous flow of H_2_ and N_2_ gas at 5 ml min^-1^ each by two thermal mass flow meters (Bronkhorst el flow, Netherlands) to maintain anaerobic conditions and provide a metabolic energy source. The outflow gas effluent was condensed using cold water from a cryostat (Julabo F25, switzerland) to prevent evaporation of the culture medium. The pH was continuously measured by a probe (Mettler Toledo InPro 3253, Switzerland) and maintained at 10.5 ± 0.5 by automatic addition of 1M NaOH with a peristaltic pump. All equipment was connected to a Wago PLC (France) via a serial link (RS232/RS485), 4–20 mA analog loop or a digital signal. The PLC was connected to a computer for process monitoring and data acquisition (BatchPro software, Decobecq Automatismes, France).

Four culture conditions were inoculated with 35 ± 5 g of chimney slurry and supplied with 5 mM of sodium acetate, sodium formate, glycine, or sodium bicarbonate as the sole carbon source. The cultures were performed in batch mode using 1L of minimal growth medium, which was inspired by the natural composition of Prony Bay hydrothermal fluid [21]: 3.7 mM NH_4_Cl, 0.92 mM K_2_HPO_4_, 86 mM NaCl, 12 mM MgCl_2_(6H_2_O), 5 mM Na_2_S_2_O_3_(5H_2_O), 5 mM Na_2_SO_4_, 3 mM CaCl_2_(2H_2_O), 5 mL L^-1^ Balch’s trace element solution [27], 0.4 mL L^-1^ Balch’s vitamin solution [27] and 1 mL L^-1^ trace metal solution. The latter contained 1.42 g L^-1^ FeSO_4_, 1.6 g L^-1^ NiSO_4_ (6H_2_O), 38 mg L^-1^ Na_2_WO_4_(2H_2_O) and 3 mg L^-1^ Na_2_SeO_3_(5H_2_O). Sulfate and thiosulfate were incorporated in the growth media as electron acceptors (thiosulfate can also be used as an energy source).

Each culture condition was tested twice: the first experimental run included the cultures Ac1, Fo1, G1, Bc1 and the second experimental run the cultures Ac2, Fo2, G2, Bc2. All cultures were maintained for 25 or 26 days and sampled every 2-3 days. Samples were immediately subdivided and prepared for different types of analysis.

### Biochemical analysis

The consumption of acetate and formate was monitored via high-performance liquid chromatography. Samples were centrifuged at 9000 rpm for 10 min at 4°C. Subsequently, 20 μL of the recovered non-diluted supernatant was injected into the column at 35°C (Dionex Ultimate 3000 from Thermofisher Scientific, USA) and eluted at 0.6 mL min^-1^ with a solution of 2.5 mM H_2_SO_4_. In addition, the consumption of sulfate and thiosulfate as electron acceptors was measured via ion chromatography. 10 μL of 1:10 diluted supernatant was injected into the column (Dionex AS-DV from Thermofisher Scientific, USA) and eluted at 1 mL min^-1^ with a 9 mM buffer solution of Na_2_CO_3_ and NaHCO_3_.

### Culture growth

The number of cells was monitored via fluorescence microscopy. Cells were fixed with formaldehyde (final concentration in sample 2%) and diluted with sterile growth media according to the cellular density. Subsequently, 2 ml of diluted sample was filtered onto white polycarbonate filters (0.2 μm porosity and 25 mm diameter from Nucleopore Whatman, USA), which were then rinsed with sterile growth medium using a vacuum pump. After filtration, the polycarbonate filters were placed on microscope slides for staining with 10-20 μL of 3.15 µM DAPI (diluted in mounting solution from CitiFluorTM AF1 Electron Microscopy Sciences, USA). Cells were counted in 15 random fields using an OLYMPUS BX-61 microscope with a filter at 372 nm (Evident, Japan). Samples were analyzed in duplicates.

### Alpha and beta diversity

The taxonomic composition at three different time points per culture (on the first day, in the middle, and on the last day), plus the inoculum of each experimental run, was assessed via 16S rRNA analysis. Samples were centrifuged at 7000 rpm and 4°C for 30 min. The pellet was recovered, and the DNA was extracted following the standard protocol of the ZymoBIOMICS DNA/RNA Miniprep Kit (Zymo Research, USA). The yielded DNA quantity and quality were evaluated on a NanoDrop One (Thermofisher Scientific, USA) and a Qubit 2.0 Fluorometer (Thermofisher Scientific, USA). Amplification and Illumina MiSeq sequencing of the 16S rRNA V3-V4 region was performed on the sequencing platform of MrDNA (Texas, USA) with two primer pairs targeting prokaryotes (341F: 5-CCTACGGGNBGCWSCAG and 805R: 5-GACTACNVGGGTATCTAATCC) and archaea (344F: 5-AYGGGGYGCASCAGGSG and 806R: 5-GGACTACVSGGGTMTCTAAT) respectively.

The bioinformatic analysis was performed in R and separately for each primer pair. Filtering and trimming of demultiplexed reads were performed in DADA2 [28], and samples with a low number of reads (< 180) were removed (one intermediate sample from the culture grown on glycine for prokaryotic sequences and one final sample each for the cultures grown on acetate, formate, bicarbonate and glycine for archaeal sequences). Denoised and dereplicated reads were merged into ASVs, allowing no mismatches in the overlapping region of forward and reverse reads. Taxonomic assignment of ASVs was performed based on the SILVA database version 138.1 [29] with a minimum bootstrap value of 60. The prokaryotic data set was rarefied to the minimum number of reads (11420) (Supplementary Figure 2A). The archaeal data set contained many bacterial ASVs, which were pruned to avoid bias in archaeal community analysis. Subsequently, the archaeal data set was rarefied to a depth of 12% (721 reads) (Supplementary Figure 2B), preserving 88% of samples left after the initial quality control in DADA2 [28]. Alpha diversity indices were calculated using the microbiome package [30] and the Microbiota Process package [31]. Subsequently, a centered log-ratio transformation was performed on the read counts using the zCompositions package [32]. On the transformed data, an Aitchison distance matrix (k = 5) was constructed for both prokaryotic (Supplementary Figure 3A) and archaeal (Supplementary Figure 3B) sequences via non-metric multidimensional scaling (NMDS) using the vegan package [33]. Permutational multivariate analysis of variance (PERMANOVA) tests were conducted to test the effect of carbon source, sampling timepoint, and experimental run on the observed sample variation. To ensure the validity of this analysis, a combined PERMANOVA model was calculated on the interaction of carbon source and sampling timepoint to inspect whether the data’s time series structure significantly affected the observed sample variation and whether it was reasonable to assess sample variability with separate models. Furthermore, analysis of variance (ANOVA) tests was performed to verify if observed differences in distance were confounded by differences in dispersion amongst groups. To identify which ASVs were driving the individual variables’ effects on sample variation the most, coefficients for the respective variable groups were extracted from the PERMANOVA model using the adonis function of the vegan package [33]. ASV relative abundance was calculated with the phyloseq package by normalizing ASV read counts to total sample read counts [34].

## Results

Cellular net growth was higher in the microbial consortia grown on acetate (the log mean number of cells mL^-1^ was 9.31E+06 in Ac1 and 1.53E+06 in Ac2) than formate (3.27E+06 in Fo1 and 7.53E+05 in Fo2), bicarbonate (1.57E+06 in Bc1 and 4.59E+05 in Bc2) and glycine which did not feature much growth (1.04E+06 in G1 and 1.58E+05 in G2). In the second run, net growth was reduced by an approximate factor of ten (Figure 2) and the net consumption of acetate and formate was much lower than run 1 (4.65 mM-acetate in Ac1 vs. 1.23 mM in Ac2, Overall, the number of bacterial taxa (853 ASVs before and 838 ASVs after rarefaction) was much higher than that of archaeal taxa (71 ASVs before and 66 ASVs after rarefaction). Prokaryotic alpha diversity and evenness of prokaryotic taxa were higher in the inoculum samples than the microbial consortia grown on glycine; followed by the consortia grown on formate, bicarbonate, and acetate (Figure 3A, Supplementary Table 1A). Archaeal alpha diversity was higher in the microbial consortia grown on glycine, than the inoculum; it did not differ much between the consortia grown on formate, bicarbonate, and acetate. Evenness of archaeal taxa was higher in the inoculum than in the consortia grown on glycine, followed by the consortia grown on formate, acetate and bicarbonate (Figure 3B, Supplementary Table 1B). Beta diversity showed considerable overlap between prokaryotes (Figure 4A) and archaea (Figure 4B) of the different microbial consortia. Nevertheless, significant differences in prokaryotic variation were observed by NMDS. According to PERMANOVA tests, the carbon source explained 29.12 % of this variation (F = 2.053, p = 0.009), the time of sampling 22.3 % (F = 2.01, p = 0.023), and the experimental run number 14.45 % (F = 1.859, p = 0.048). The interaction between carbon source and sampling time was insignificant (F = 0.25, p = 0.154), with the individual effect of both variables remaining significant in the combined model, validating the statistical model used. Two *Rhodobacteraceae* ASVs (one classified to the order level as *Roseibaca*) were the most significant drivers of sample dissimilarity by carbon source, indicated by their relatively high PERMANOVA coefficients (ASV_289_ = 2.459, ASV_004_ = 2.212) (Figure 5A). For the sampling time, a *Meiothermus* ASV was the most significant driver of sample dissimilarity (ASV_003_ = 3.208) (Figure 5B). For the experimental run number, the most critical drivers of sample dissimilarity were two ASVs associated with *Pseudahrensia* and *Dethiobacteraceae,* respectively (ASV_345_ = 4.325, ASV_119_ = 4.286) (Figure 5C). ASVs driving sample similarity in both carbon source and sample time were associated with the cluster of uncultivated MSBL5 *(Dehalococcoidia)* (carbon source: ASV_042_ = -2.76, sampling time: ASV_006_ = -3.99), *Acetothermia* (carbon source: ASV_011_ = -2.4, sampling timepoint: ASV_005_ = -3.09 and ASV_011_ = -3.23)*, Desulfovibrio* (carbon source: ASV_016_ = -2.09, sampling timepoint: ASV_016_ = -2.78) and ML635J-40 aquatic group *(Bacteroidales)* (carbon source: ASV_007_ = -2.11, sampling timepoint: ASV_007_ = -2.95) taxa (Figure 5A,B). For the archaea, the carbon source and run number had no significant effect on the distance matrix (F = 1.229, p = 0.15 with R^2^ = 0.197 and F = 1.183, p = 0.26 with R^2^ = 0.097 respectively). Sampling time slightly affects community composition, explaining only 8.2 % of the variance (F = 1.558, p = 0.053). The most significant driver of sample dissimilarity according to the sampling time was an ASV identified as *Woesearchaeales* (ASV_037_ = 1.01). Drivers of sample similarity were ANME-3 (ASV_007_ = -2.93, ASV_019_ = -2.88, ASV_003_ = -1.4, ASV_039_ = -1.37, ASV_045_ = -1.36, ASV_002_ = -1.24) and *Syntrophoarchaeaceae* (ASV_006_ = -4.33, ASV_034_ = -1.76, ASV_061_ = -1.23) which are both associated with the *Methanosarcinales* (Figure 5D). All ANOVA tests were non-significant, confirming the reliability of the variables’ effects on observed differences in prokaryotic and archaeal distance matrices.

**Figure 2.**
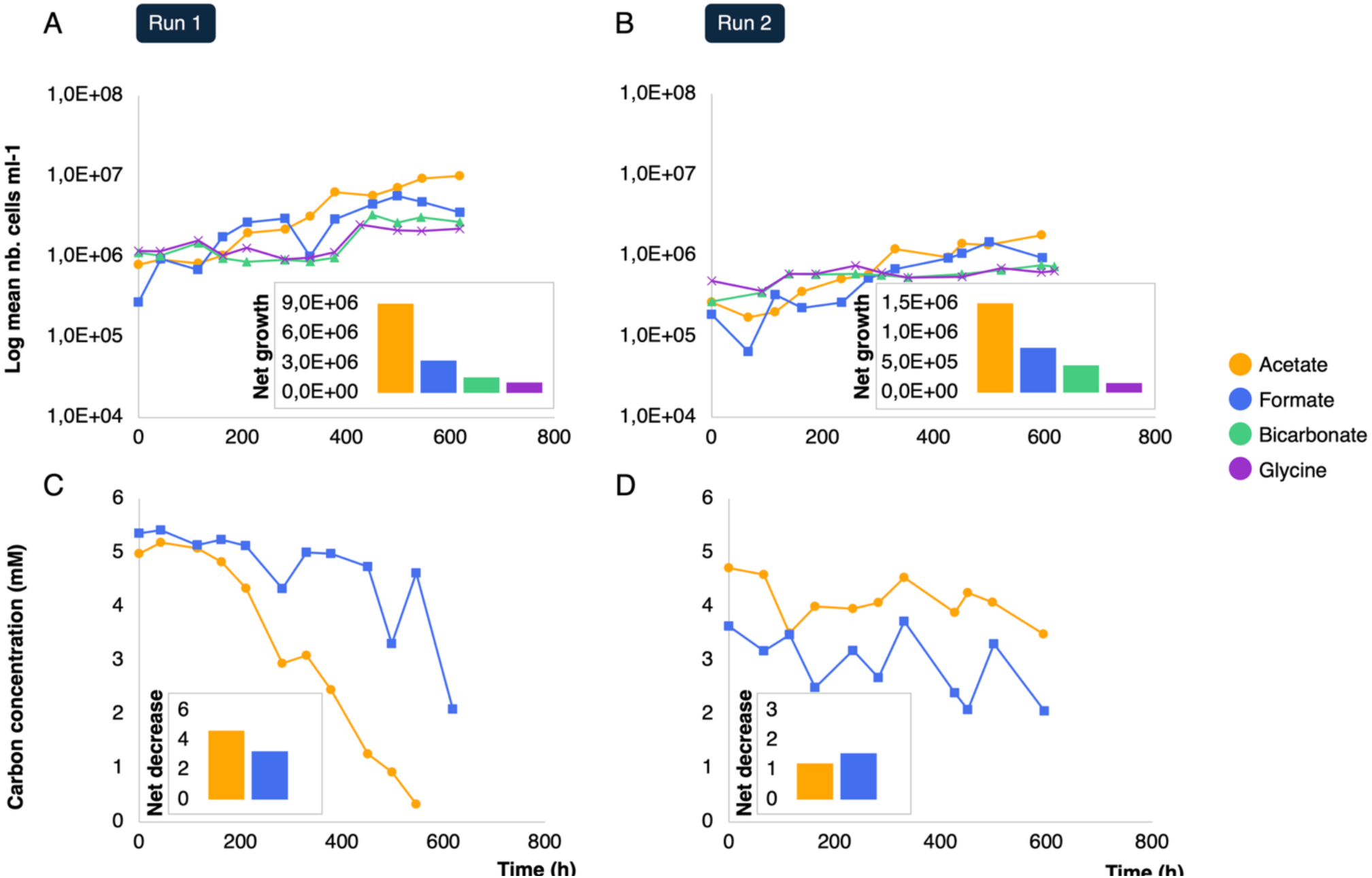
Microbial growth and carbon consumption in the first experimental run (A, C; cultures Ac1, Fo1, G1, Bc1) and in the second run (B, D; cultures Ac2, Fo2, G2, Bc2). The number of cells was counted via DAPI-staining and fluorescence microscopy. The concentration of acetate and formate in the respective consortia was determined by high-performance liquid chromatography. The net increase of cells and the net decrease of acetate and formate (shown as inserts) were calculated as the difference between the first and last sampling timepoint.

**Figure 3.**
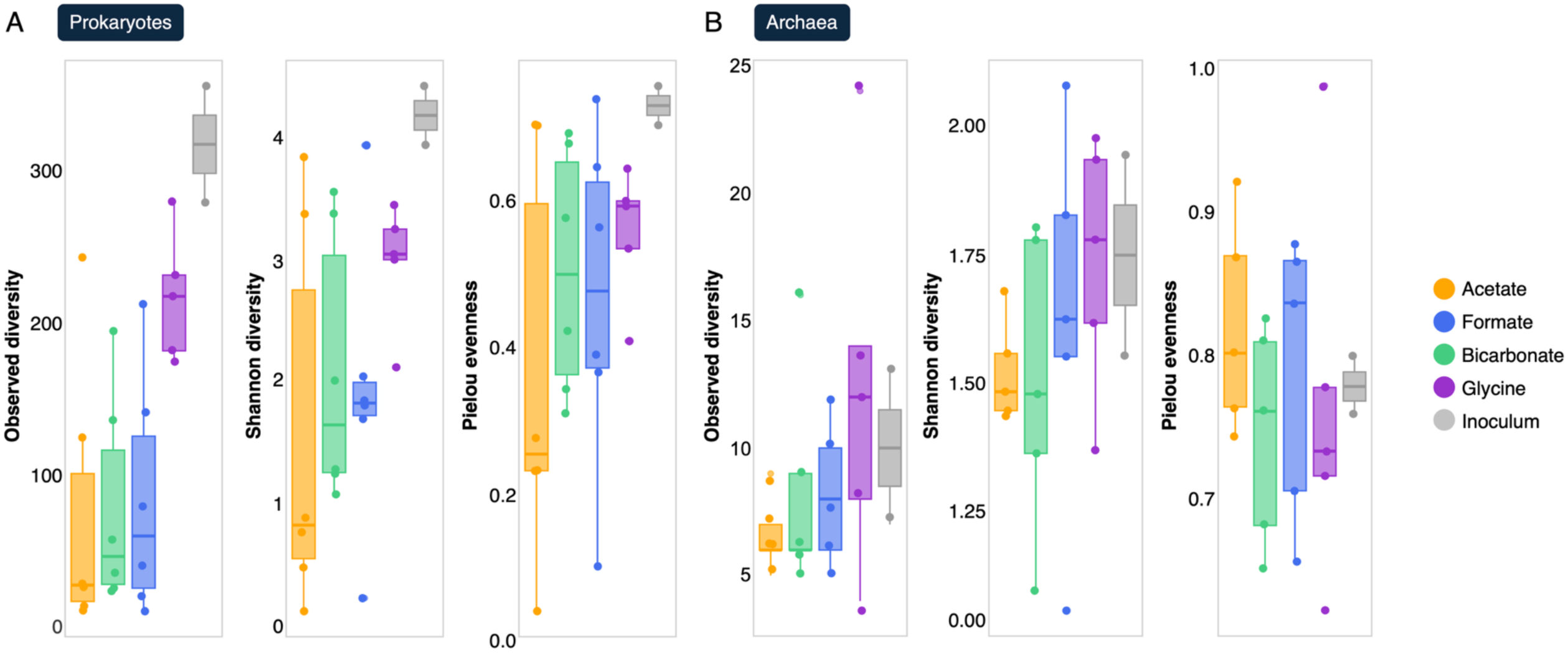
Alpha diversity and evenness calculated to the observed number of ASVs, the Shannon index and the Pielou index in prokaryotic (A) and archaeal (B) sequences from the inoculum (grey) and microbial consortia grown on acetate (yellow), formate (blue), bicarbonate (green) or glycine (purple). Values for the same conditions from both experimental runs are combined.

**Figure 4.**
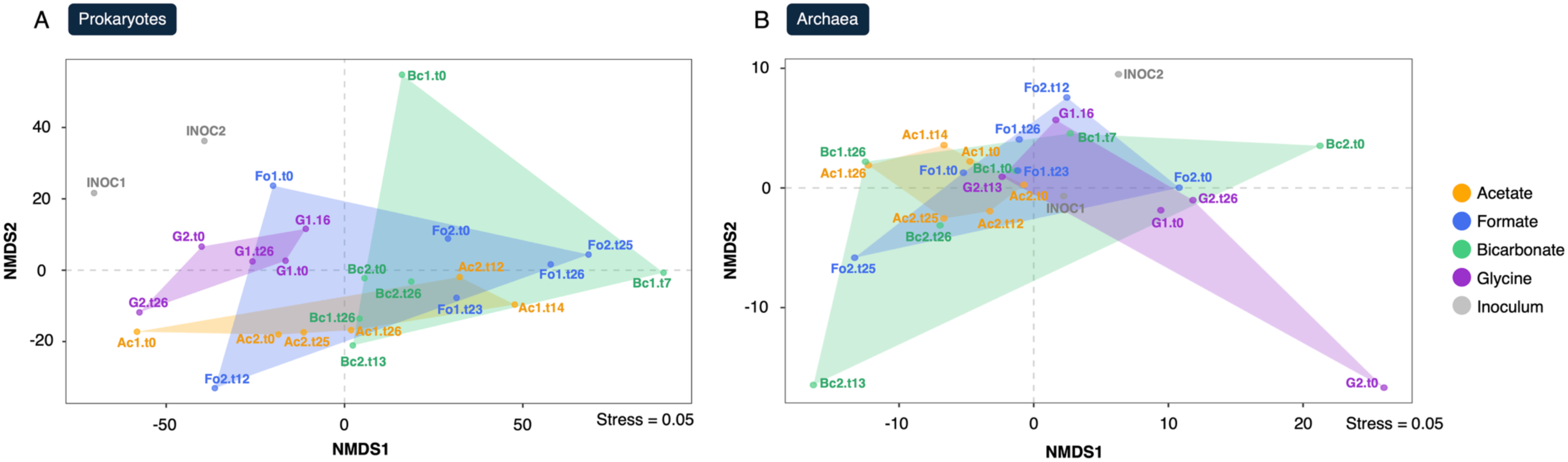
Aitchison distance matrices (k = 5) constructed on prokaryotic (A) and archaeal (B) sequences from the inoculum (grey) and microbial consortia grown on acetate (yellow), formate (blue), bicarbonate (blue) or glycine (purple). Distance matrices were constructed using a non-metric multidimensional scaling on log-ratio transformed data. The name of cultures Ac1, Fo1, G1, Bc1 (corresponding to the run 1) and cultures Ac2, Fo2, G2, Bc2 (corresponding to the run 2) also indicated the sampling time (t0, t7, t12, t13, t16, t23, t25, t26).

**Figure 5.**
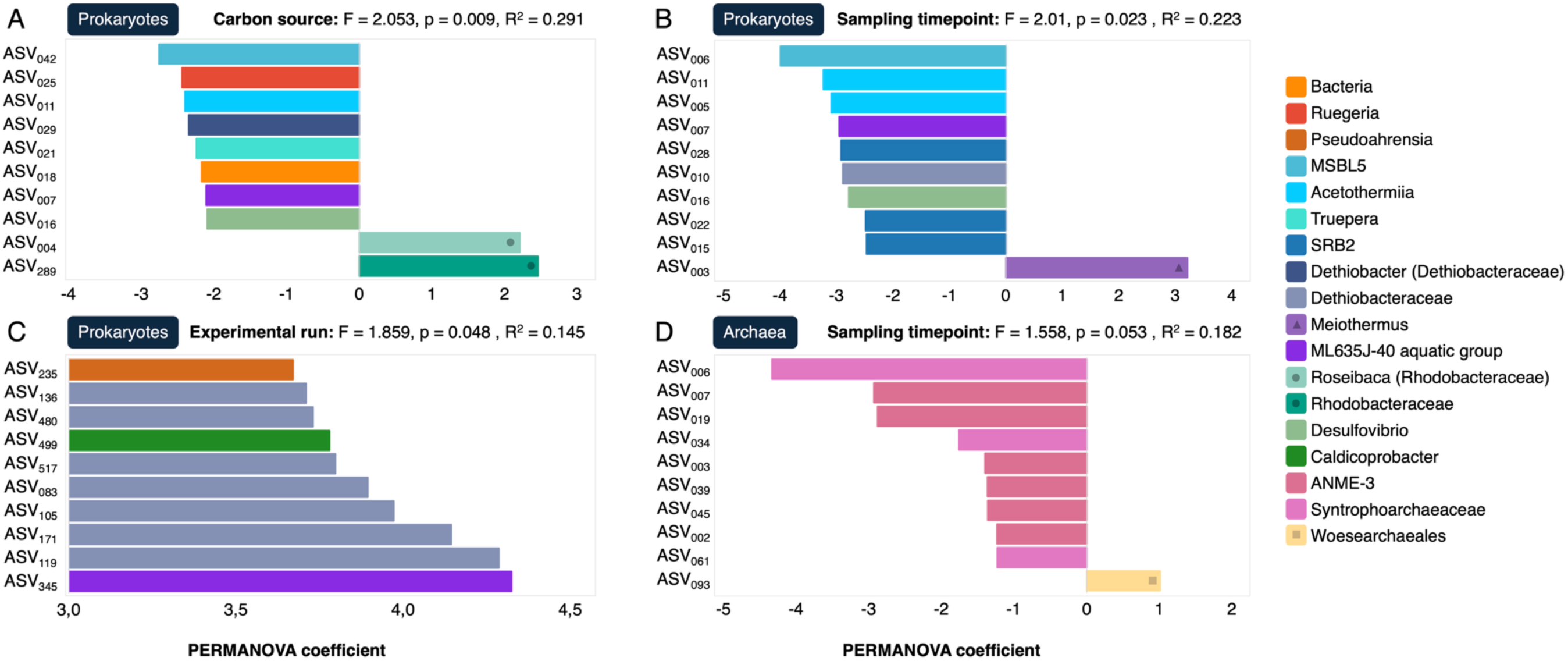
Top ASVs driving significant effects of carbon source, sampling timepoint and experimental run on sample variation in prokaryotic (A, B, C) and/or archaeal (D) sequences. Different ASVs are color-coded according to the lowest taxonomic level to which they were assigned. Coefficients for the respective group of variable group were calculated in permutational multivariate analysis of variance tests; corresponding F values, p values and R^2^ value are given on the top right of each graph. Positive coefficients indicate an effect of the respective ASV on sample dissimilarity, negative coefficients indicate an effect on sample similarity. Key taxonomic groups are highlighted with symbols to improve the readability of the graph.

The evolution of the taxonomic composition over time in the different microbial consortia was consistent with alpha and beta diversity predictions. On all carbon sources except glycine, the prokaryotic community changed significantly over time, and enrichment of specific taxonomic groups was observed (Figure 6A). The inoculum and samples from the first day of culture were mainly composed of the genera *Acetoanaerobium* and *Roseibaca* and a relatively smaller proportion of *Methyloceanibacter*, *Truepera*, *Desulfovibrio*, *Caldicobacter*, *Filomicrobium,* and *Ruegeria*. Acetate-grown cultures displayed a large proportion of *Roseibaca* and a small proportion of *Halomonas* after 12 (Ac2) and 14 (Ac1) days, respectively. The strong enrichment of *Roseibaca* was maintained until the end of the culture, next to a lesser enrichment of *Meiothermus* (Ac1 and Ac2). Formate-grown cultures also showed a strong enrichment of *Roseibaca* (Fo1 and Fo2) and a slighter enrichment of *Meiothermus* (Fo1) and *Thioalkalimicrobium* (Fo1) at the end of experiments, and a predominance of *Acetoanaerobium* (Fo2) after 12 days. On bicarbonate, the opposite was observed: a strong enrichment of *Meiothermus*, and a less pronounced (re-)enrichment of *Roseibaca* at the end of the culture (Bc1 and Bc2), next to an enrichment of *Hydrogenophaga* (exclusively observed in Bc1). On day 7, the culture was dominated by *Halomonas* next to *Thioalkalimicrobium* and *Reinekea* enrichments (Bc1), and on day 13, *Roseibaca* next to a slight enrichment of *Halomonas* (Bc2). Little development was observed on glycine: the community structure resembled the inoculum throughout the experiment in both runs 1 and 2. The archaeal community (Figure 6B) was strongly dominated by the order *Methanosarcinales* (ASV classified to the genus level as ANME-3) in the inoculum, next to a small proportion of *Nitrosopumilales* (INOC2). They were also predominant in all culture samples except for the end of the culture grown on bicarbonate, where a strong enrichment of *Methanomicrobiales* was observed (Bc2). In addition, a slight enrichment of *Woesearchaeales* was observed in the middle and at the end of the culture grown on acetate (Ac1 and Ac2) and formate (Fo1), as well as on day 7 of the culture grown on bicarbonate (Bc1). They were also present at the beginning (G2), middle, and end (G1) of the culture grown on glycine, but their relative abundance decreased over time.

**Figure 6.**
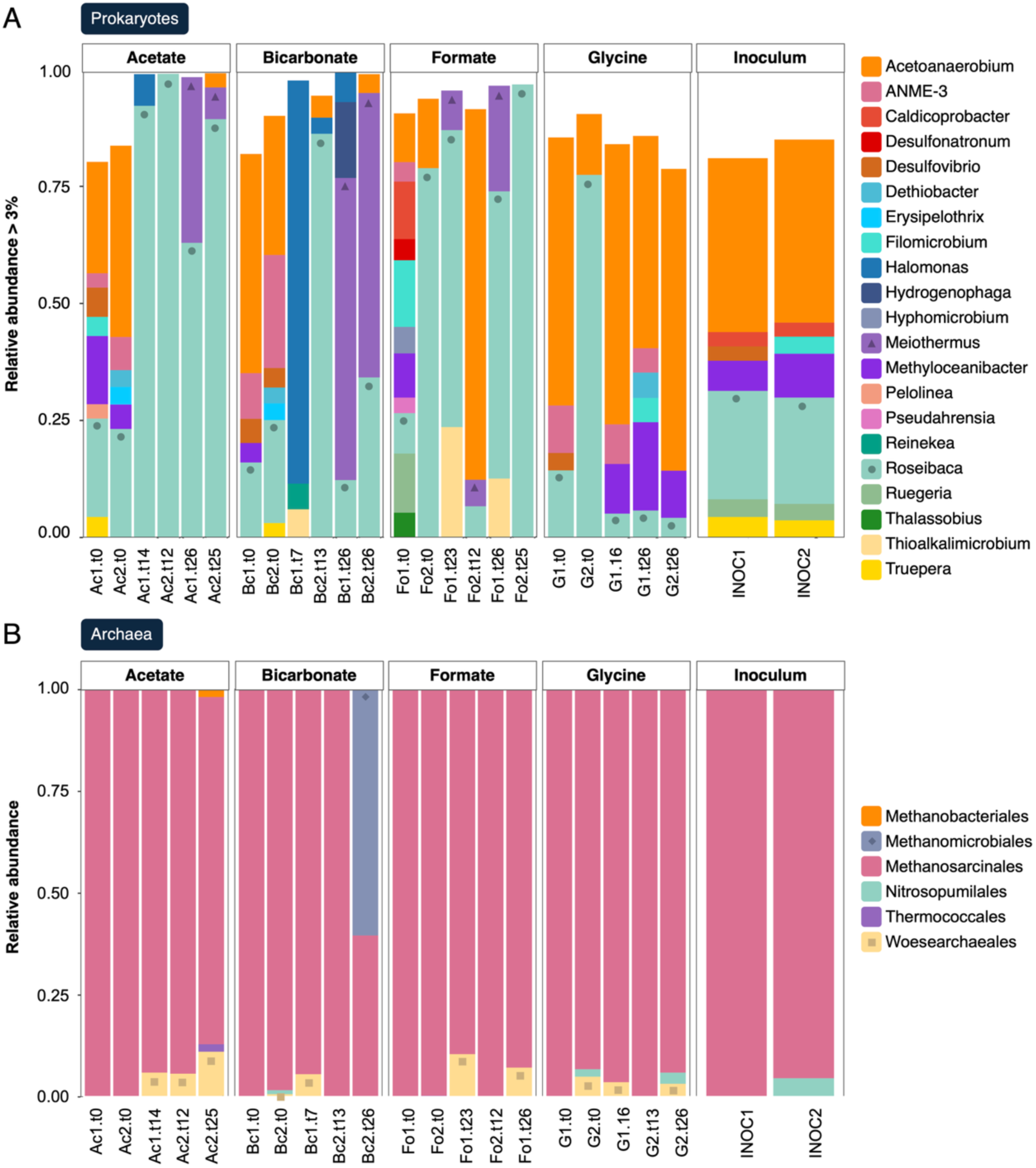
Taxonomic composition of prokaryotic (A) and archaeal (B) microbial consortia grown n acetate (Ac), bicarbonate (Bc), formate (Fo) or glycine (G) in the first (1) and second (2) xperimental run over time. For each carbon source and experimental run, the taxonomic omposition of up to three different timepoints is depicted, including the first day (t0), an ntermediate day and the last day (t25 or t26) of the culture. In addition, the taxonomic composition f the inoculum used in the respective experimental run (INOC) is shown. For bacterial sequences, nly genera with a relative abundance > 3% are depicted. For archaeal sequences, all orders are hown. Note that ANME-3 is a genus of the archaeal order *Methanosarcinales* which was also mplified by the prokaryotic primers. Key taxonomic groups are emphasized with symbols to mprove readability of the graph.

## Discussion

### The natural community

The natural prokaryotic community was dominated by several taxonomic groups that are commonly found in Prony Bay and other serpentinite-hosted environments. *Acetoanaerobium* is an anaerobic bacterium that can produce acetate from the fermentation of complex carbohydrates; it is one of the strains that has been isolated from Prony Bay hydrothermal chimneys [35]. Close relatives of *Desulfovibrio*, an anaerobic sulfate-reducing bacterium, have been found in Prony Bay, at the serpentinized Costa Rica Margin and in Lost City [14, 23, 36]. Likewise, the facultatively anaerobic chemoorganotroph *Truepera* has been documented in Prony Bay and in the Coast Range ophiolite [24, 37]. None of these marker taxa were significantly enriched in our culture conditions (except *Acetoanaerobium* on formate), suggesting that they rely on the metabolic products of other community members or that their growth is limited by other experimental conditions such as light.

### Carbon sources of cultivated microbial consortia

All carbon sources facilitated culture growth and development except for glycine. The microbial consortium maintained on glycine featured a very low growth rate (Figure 2), and diversity and evenness scores remained consistently elevated (Figure 3), suggesting no significant enrichment of taxonomic groups specialized on the uptake of this compound. This was confirmed by the taxonomic composition of the consortia, which did not change much over time and remained similar to the natural community (Figure 6). The use of glycine as a carbon source in a serpentinization context was proposed by Nobu et al. [10], who showed that glycine reductase genes are abundant in metagenomes from various serpentinite-hosted systems. The glycine reductase complex is a central enzyme in the reductive glycine pathway, a recently proposed seventh carbon fixation pathway [16]. While it is indeed possible for exogenous glycine to enter the glycine reductase complex via a membrane-bound transporter, it might also be produced as an endogenous intermediate in the reductive glycine pathway. Similar to the other carbon fixation pathways, the reductive glycine pathway starts with one molecule of CO_2_ that could be yielded from both formate [13] and bicarbonate [13, 14]. It would be therefore the primary carbon source in this case. It is likely that in Prony Bay, the use of formate and bicarbonate as carbon sources is more favorable compared to glycine whose natural concentration is expected to be low [10]. This might explain the low growth observed in this experimental condition.

Contrary to glycine, microbial growth was observed in cultures grown in presence of bicarbonate, formate and especially acetate (Figure 2). This was accompanied by a steep decrease in diversity and evenness (Figure 3), suggesting the enrichment of specific taxonomic groups consuming these carbon sources. This is confirmed by the evolution of the taxonomic composition of the cultures after the second and third week of incubation (Figure 6). The main driver of sample dissimilarity according to carbon source was the *Roseibacaceae* family (Figure 4, Figure 5).

### Dominant taxa of cultivated microbial consortia

The consortia grown on acetate and formate were heavily dominated by the genus *Roseibaca* previously detected in the Aqua de Ney serpentinizing spring (Trutschel et al., 2022, 2023). Members of this genus are anoxygenic photoheterotrophs and rose-colored due to their bacteriochlorophyll pigments [38, 39]. They have previously been detected in Prony Bay, where their biofilms may be responsible for the dark-pink color of the chimney exteriors [24]. As all our cultures were incubated at O_2_ concentrations strictly below 0.5%, the growth of *Roseibaca*, characterized as strict aerobes, was quite unexpected (Supplementary Figure 4). Since the experiments were performed in clear glass bioreactors, ambient light could have favored the growth of oxygenic phototrophs locally supplying O_2_. However, no significant occurrence of cyanobacteria (the only bacterial group that performs oxygenic photosynthesis) was observed [40] (Figure 6). The cultivated *Roseibaca* members may therefore be facultative anaerobes. The carbonaceous chimney wall is very porous, and at a depth where O_2_ becomes scarce, light might still penetrate. The bacteriochlorophyll pigments used in anoxygenic photosynthesis can absorb light at near-infrared wavelengths and facilitate photosynthesis in deep water or sediments [40]. In addition, the occurrence of normally strictly aerobic taxa in anoxic serpentinized fluids is a relatively common phenomenon and has been suggested to drive localized speciation of closely related populations along the steep O_2_ and redox gradients from surface waters to subsurface fluids [41].

Next to *Roseibaca*, the consortia grown on acetate and formate included a significant proportion of *Meiothermus,* previously detected in serpentinizing springs of New Caledonia [42, 43] in the Zambales Ophiolites, Philippines [44], and in the Samail Ophiolite, Oman [11] The cultures alimented with formate additionally featured a slight enrichment of *Thioalkalimicrobium*, reclassified into *Thiomicrospira* [45]. *Thioalkalimicrobium* or *Thiomicrospira* were previously observed in submarine serpentinizing Lost City and Prony bay hydrothermal systems (Brazelton et al., 2010, 2012; Postec et al., 2015) and terrestrial serpentinizing Aqua de Ney spring (Trutschel et al., 2023). They were related to the thiosulfate consumption observed in formate-grown cultures (see Supplementary Fig. S1) and are most likely thiosulfate-oxidizing bacteria and also known as H_2_-oxidizers [46]. The formate-grown cultures also displayed mid-term enrichment of *Acetoanaerobium,* an anaerobic genus previously isolated from Prony Bay (Bes et al., 2015). The latter probably fermented organic compounds produced by the consortium during the first week and does not count amongst primary producers.

The consortia grown on bicarbonate was strongly dominated by the *Meiothermus* genus. Similar to *Roseibaca*, *Meiothermus* has been described as a strictly aerobic heterotroph; however, Munro-Ehrlich et al. [41] provided evidence that subsurface populations can undergo parapatric speciation and develop traits of facultatively anaerobic autotrophs, such as the gain of [NiFe] hydrogenases. Similar to these subsurface genotypes, it is likely that the *Meiothermus* population cultivated on bicarbonate grew chemoautotrophically by producing CO_2_ in the cytoplasm with H_2_ as electron donor. Besides *Meiothermus,* a significant enrichment of the chemoautotrophic *Hydrogenophaga,* a sister group of H_2_-oxidizing *Serpentinimonas* (a marker genus of serpentinization) which features bicarbonate-uptake genes, could be observed along with enrichment of *Roseibaca* [4, 47]. While *Roseibaca* are usually classified as photoheterotrophs, their consistent and significant growth on bicarbonate might suggest that we enriched a novel photoautotrophic genotype. In addition, the genus *Halomonas* was dominant in the second week of the first experimental run, a thiosulfate-oxidizing bacterium that is typically observed in serpentinite-hosted environments. As this genus is heterotrophic, however, it is unlikely that it used bicarbonate as a carbon source [48–50] (Figure 6).

The initial archaeal community consisted almost exclusively of the ANME-3 cluster, which belongs to the *Methanosarcinales*. This group was not enriched but maintained throughout the entirety of the cultivation experiments on all carbon sources, with a predominance so strong that the taxonomic composition did not vary significantly between carbon sources (Figure 6B). The *Methanosarcinales* are one of the most typical taxa associated with serpentinization-hosted ecosystems. Two phylotypes belonging to this group have been described. The Lost City *Methanosarcinales* (LCMS) are more closely related to ANME-3 (our most abundant ANME-3 ASV shares 95.31% sequence identity with the LCMS), while The Cedars *Methanosarcinales* (TCMS) are intermediate between the ANME-2c and *Methanomicrobiales* clusters [51, 52]. While ANME-3 are classified as anaerobic methane oxidizers (AOM), it has been shown that they possess all the genes necessary for methanogenesis and LCMS are usually described as methanogens. It is, therefore, unclear whether LCMS are consumers or producers of methane, or both [51]. In addition to the maintenance of *Methanosarcinales*, a slight enrichment of *Woesearchaeales* was observed on formate, bicarbonate and particularly acetate in the first experimental run. This enrichment accounted for an almost significant influence of the sampling time on the community structure (Figure 5D). Since the abundances shown here are relative, the archaeal taxonomic composition may reflect an emerging symbiotic relationship between *Methanosarcinales* and *Woesearchaeales*. The *Woesearchaeales* are probably incapable of an independent lifestyle and have been suggested to establish symbiotic relationships with methanogens [53]. So far, only heterotrophic genotypes are known [53]. While only a handful of taxa has been shown to perform SAO, it might be possible that our *Woesearchaeales* ASV is able to oxidize acetate to CO_2_ and H_2_ in presence of a *Methanosarcinales* as syntrophic partner. In the second experimental run, a remarkable enrichment of *Methanomicrobiales* was observed on bicarbonate towards the end of the culture (81.44% ASV sequence identity with TCMS). This order is methanogenic [54] and probably outcompeted the symbiosis between *Methanosarcinales* and *Woesearchaeales* (Figure 6B). Interestingly, a recent study reclassified TCMS as acetogenic *Methanocellales* featuring bicarbonate uptake genes [55]. It is thus likely that our ASV is able to fix bicarbonate as an autotrophic carbon source.

### The shallow hydrothermal chimney foundation consortium

The predominance of phototrophic *Roseibaca* in two culture conditions suggests that our results are mostly reflective of Prony Bay’s shallow hydrothermal chimney community. The pronounced growth on acetate supports the foundation species theory proposed by Lang et al. [18]. Their original model was based on a high occurrence of sulfate reducers taking up formate and releasing CO_2_ in the deep-sea Lost City hydrothermal field. While it is likely that sulfate reducers (e.g. *Desulfovibrio*, *Desulfonatronum*) play an essential role in the deeper subsurface communities of Prony Bay, they were not enriched in the bioreactors despite sulfate and thiosulfate being the sole electron acceptors added to the cultures (Figure 6, Figure 7A). On the contrary, at least two of the enriched taxa (*Thioalkalimicrobium* and *Hydrogenophaga*) have previously been reported to reduce nitrate, suggesting that the respiration of nitrate could play a role in this anaerobic natural system. The electron acceptor used by the cultivated consortia remains unclear (Figure 7A). In the bioreactors alimented with acetate, the microbial consortium of putative heterotrophic foundation species was heavily dominated by the anoxygenic photoheterotroph *Roseibaca*, suggesting that this metabolic type plays a more important role in serpentinite-hosted environments than previously assumed. Besides *Roseibaca*, the putative consortium of heterotrophic foundation species included the heterotrophic genotype of *Meiothermus* [41]. Together, *Roseibaca* and *Meiothermus* may produce CO_2_ from acetate for various autotrophs on the first trophic level in the natural ecosystem (Figure 7B).

**Figure 7.**
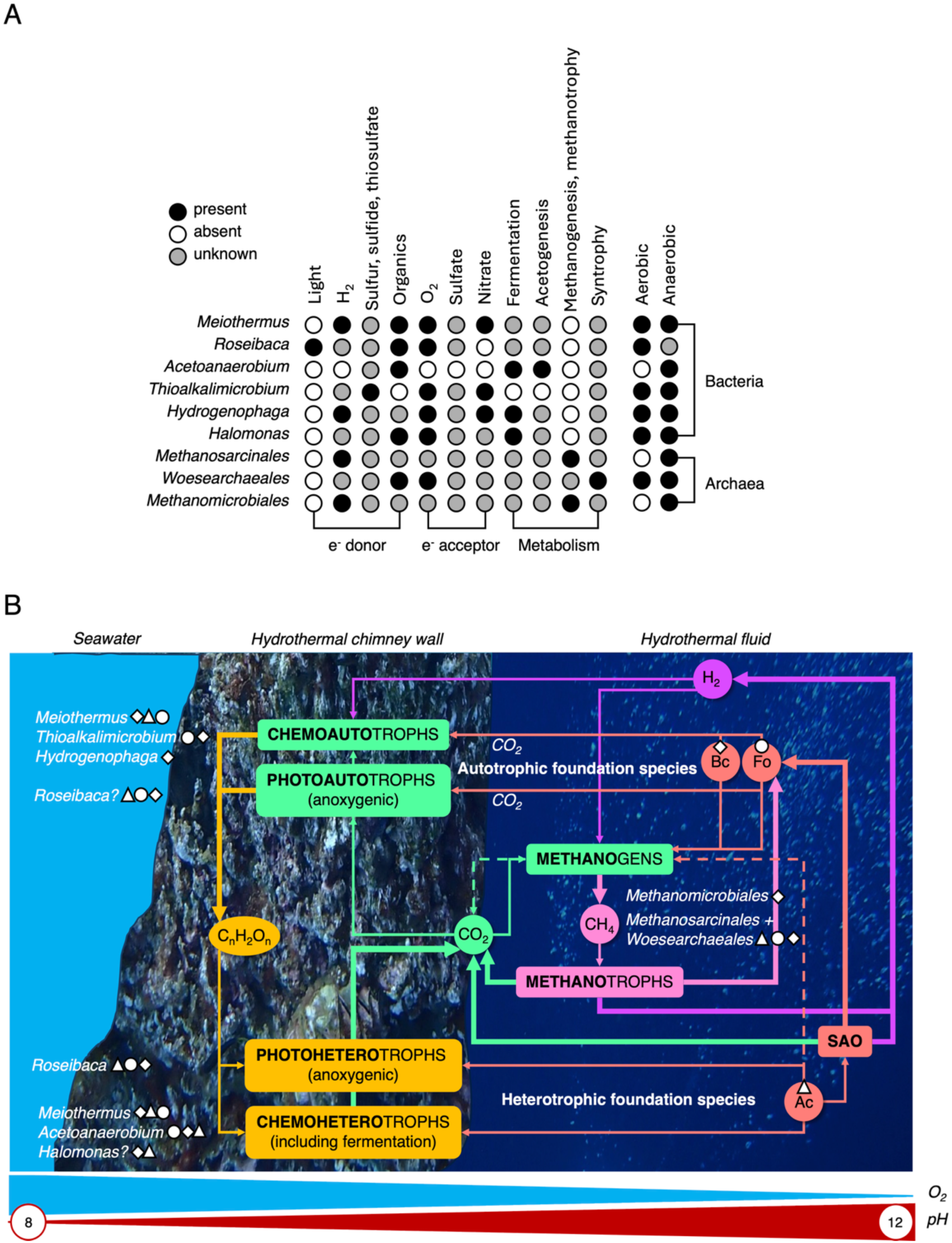
Proposed metabolic network of the cultivable Prony Bay hydrothermal community. A) Metabolic potential of enriched key taxa according to the references cited in the main text. B) Hypothesized trophic relationships. Putative foundation species are shown next to metabolic categories, with symbols indicating all culture conditions they were enriched in. Note the following abbreviations: SAO for syntrophic acetate oxidation, Bc for bicarbonate (diamond), Fo for formate (circle), Ac for acetate (triangle).

Next to these putative acetate-assimilating foundation species, we cultivated potential autotrophic foundation consortia on bicarbonate and formate. On bicarbonate, this concerned the autotrophic *Meiothermus* genotype described by Munro-Ehrlich et al. [41] and the hydrogenotrophic *Hydrogenophaga*. It is likely that the same *Meiothermus* genotype is part of base of the trophic network cultivated on formate, alongside the thiosulfate-oxidizing autotroph *Thioalkalimicrobium* and potentially a novel photoautotrophic genotype of *Roseibaca*. Together, these bicarbonate- and formate-fixing autotrophs could produce organic compounds for heterotrophs on the first trophic level such as *Acetoanaerobium* and *Halomonas* (Figure 7B).

A special case of such carbon cycling between heterotrophs and autotrophs constitutes the relationship between methanogens and methanotrophs: unidentified methanotrophs could establish a supply of CO_2_ for autotrophic methanogens and vice versa. The *Methanomicrobiales* might perform methanogenesis with bicarbonate as carbon source, thus constituting an autotrophic foundation species. The predominant *Methanosarcinales* on the other hand are likely dependent on CO_2_ that is either produced by a symbiont or locally released by other community members. While endogenous CO_2_ supply might be provided by a potentially acetate-oxidizing *Woesearchaeales*, external CO_2_ could rely on acetoclastic methanogenesis in addition to methanotrophy (Figure 7B).

## Conclusion

Specific taxonomic groups were successfully cultivated from a shallow submarine Prony Bay chimney sample in anaerobic bioreactors simulating *in-situ* conditions. These reactors were operated with artificial hydrothermal fluid under a continuous flow of H_2_ gas, supplemented with sulfate and thiosulfate as the sole electron acceptors, and bicarbonate, formate, acetate or glycine as the sole carbon source. The culture on glycine exhibited only minimal growth and the taxonomic composition barely changed compared to the initial diversity observed in the inoculum. This either suggests that glycine plays a minor role in the natural microbial ecosystem, or that the experimental conditions did not accurately reproduce the natural conditions. The latter is supported by the observation of the generally very low growth rates on glycine, which might have required longer incubation periods. The pronounced growth on acetate supports a modified version of the foundation species theory proposed by Lang et al. [18]. Heterotrophic foundation taxa such as *Roseibaca* and *Meiothermus* (heterotrophic genotype) may produce CO_2_ from acetate, facilitating the development of a diverse community of autotrophs, including those without bicarbonate and formate transporters or symbionts. Autotrophic foundation species might include bicarbonate-fixing *Meiothermus* (autotrophic genotype) and *Hydrogenophaga*, as well as formate-fixing *Meiothermus*, *Thioalkalimicrobium* and potentially a novel photoautotrophic genotype of *Roseibaca* producing organic compounds for heterotrophs such as *Acetoanaerobium* and *Halomonas.* In the archaeal community, this reciprocal relationship is illustrated by CO₂ and CH₄ cycling *Methanosarcinales* with putative acetate-oxidizing *Woesearchaeales* symbionts, as well as bicarbonate-fixing *Methanomicrobiales*. We propose that such a feedback loop contributes to the diversification of the serpentinite-hosted community and explains the presence of well-established microbial communities in environments depleted of DIC.

## Supporting information

All supplementary information

## Declarations

### Ethics approval and consent to participate

Not applicable.

### Consent for publication

Not applicable.

### Availability of data and materials

The raw 16S rRNA sequencing data analyzed in this study is available in the National Center for Biotechnology Information repository under the BioProject accession PRJNA1223160.

The sample metadata used to analyze beta diversity is included in the article and it’s additional files.

The bioinformatic pipeline developed for this study is available on github: https://github.com/jampoa/16S-alpha-beta-div

## Authors’ contributions

GE directed the project. GE and AP designed and supervised the study. GE, REP, RMP and MQ collected the samples. GE, AP, RMP, SD, REP, YCB and MQ developed the experimental design. AR, RMP, SD, and AP performed the experimental work. RMP and AR performed the initial data analysis. RMP performed the final data analysis and wrote the manuscript. All authors reviewed the manuscript and read and approved the final version.

## Acknowledgements

We are grateful to all participants of the 2022 field campaign to Prony Bay; firstly the scientific divers of the ENTROPIE & IMAGO units at the IRD Research Center Nouméa: Bertrand Bourgeois, John Butler, Magali Boussion and Mahé Dumas, without whom this mission would not have been possible; Clarisse Majorel and Isabelle Biegala for welcoming and assistance in their lab at IRD Nouméa, and Véronique Perrin for her invaluable help in resolving all administrative and logistical difficulties. We would like to thank Dr. Ariane Bize (INRAe Prose, Antony, France) for the training and discussions on DNA Stable Isotope Probing, which could not be included in this article but was greatly appreciated. Furthermore, we would like to thank the OMICS platform with Marc Garel and Fabrice Armougom (Mediterranean Institute of Oceanology UMR 7294, Marseille, France), who provided the workflow and source code that our bioinformatic pipeline is based on.

## Authors’ information

Not applicable.

