## Supplementary material for "Cultivating microbial communities from the serpentinite-hosted Prony Bay Hydrothermal Field on different carbon sources in hydrogen-fed bioreactors": All supplementary information

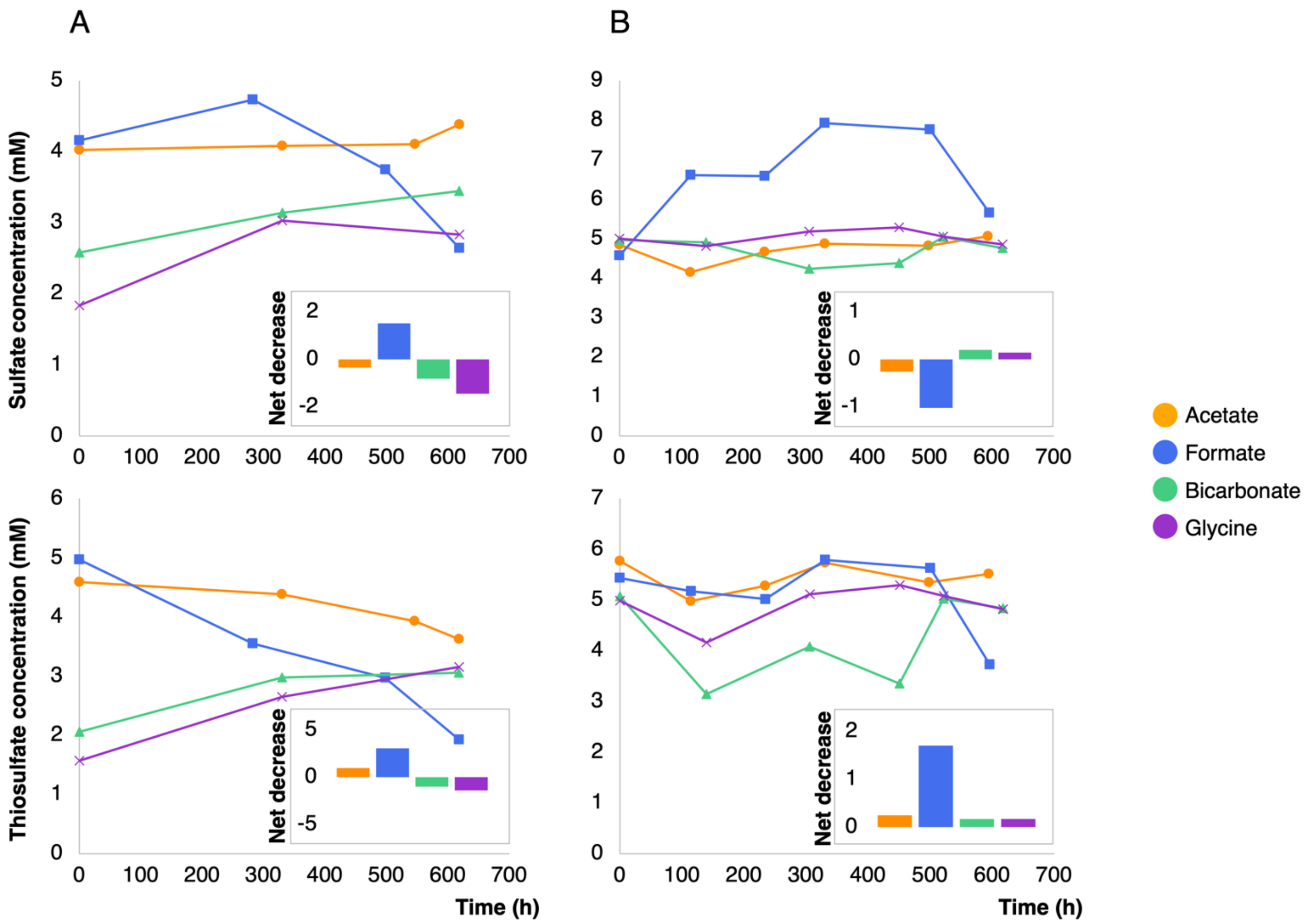


**Supplementary Figure 1.** Sulfate and thiosulfate consumption in the first (A) and second (B) experimental run in the microbial consortia grown on acetate (yellow), formate (blue), bicarbonate (green) or glycine (purple). The respective net decrease of sulfate or thiosulfate concentration, calculated as the difference between the first and last timepoint measured, is illustrated with barplots.

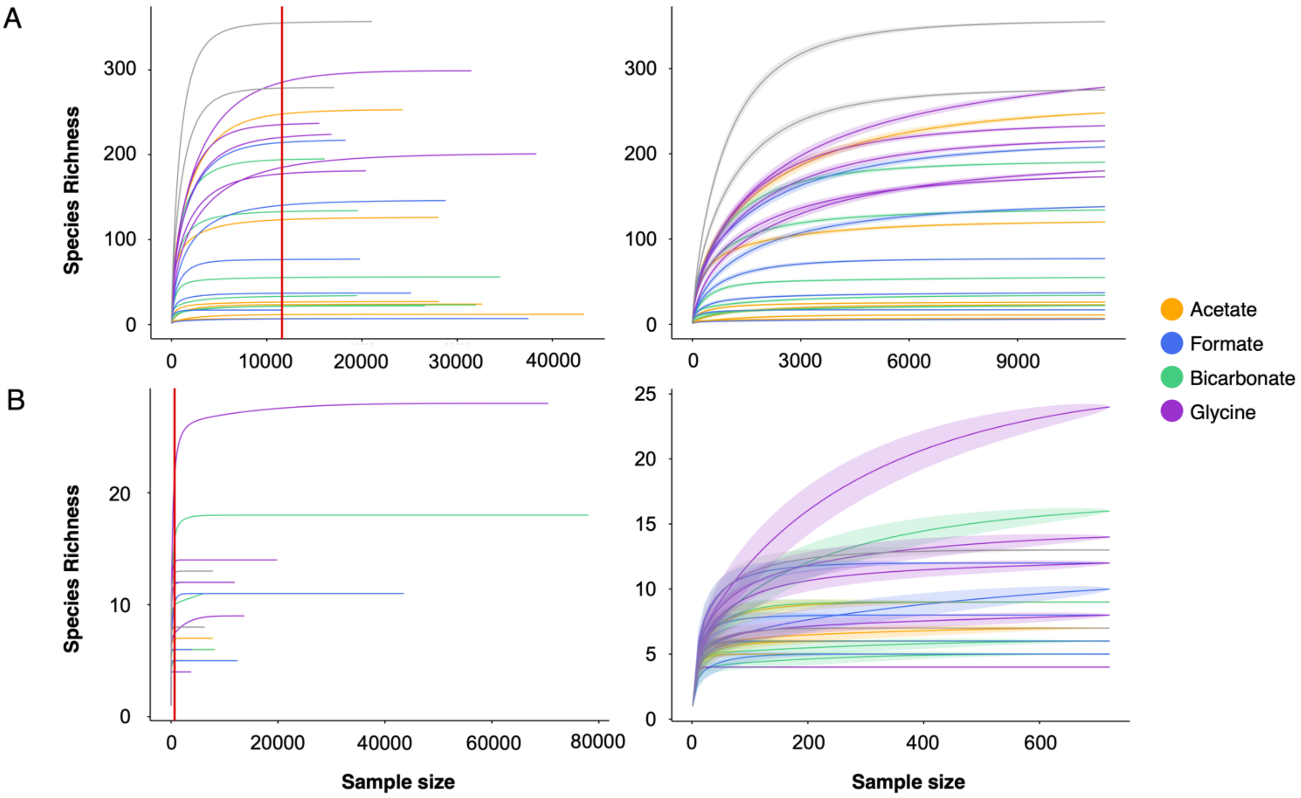


**Supplementary Figure 2.** Rarefaction curves for bacterial (A) and archaeal (B) sequences from the microbial consortia grown on acetate (yellow), formate (blue), bicarbonate (green) or glycine (purple). Bacterial sequences were normalized to 11420 reads, which corresponds to the minimal sample size, archaeal sequences to 721 reads. The rarefaction depth is indicated with a red vertical line.

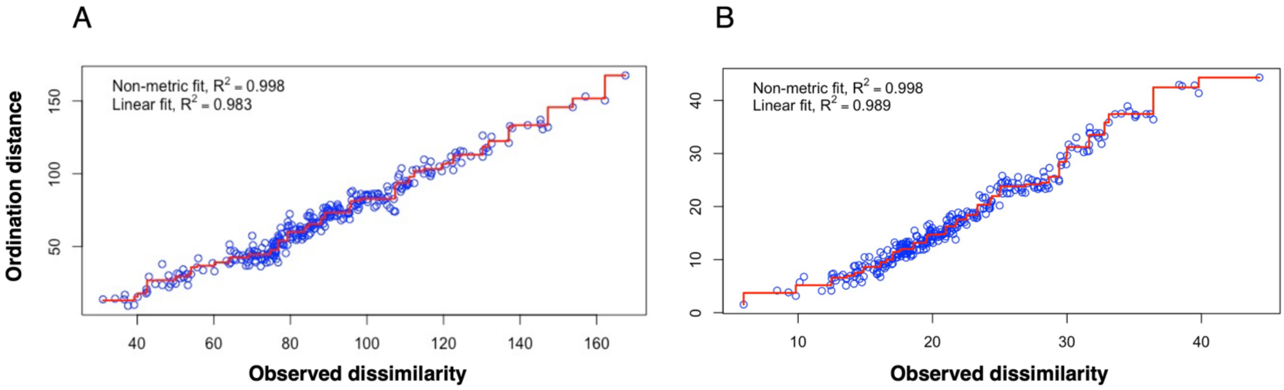


**Supplementary Figure 3.** Stressplots of the non-metric multidimensional scaling matrices calculated for bacterial (A) and archaeal (B) sequences.

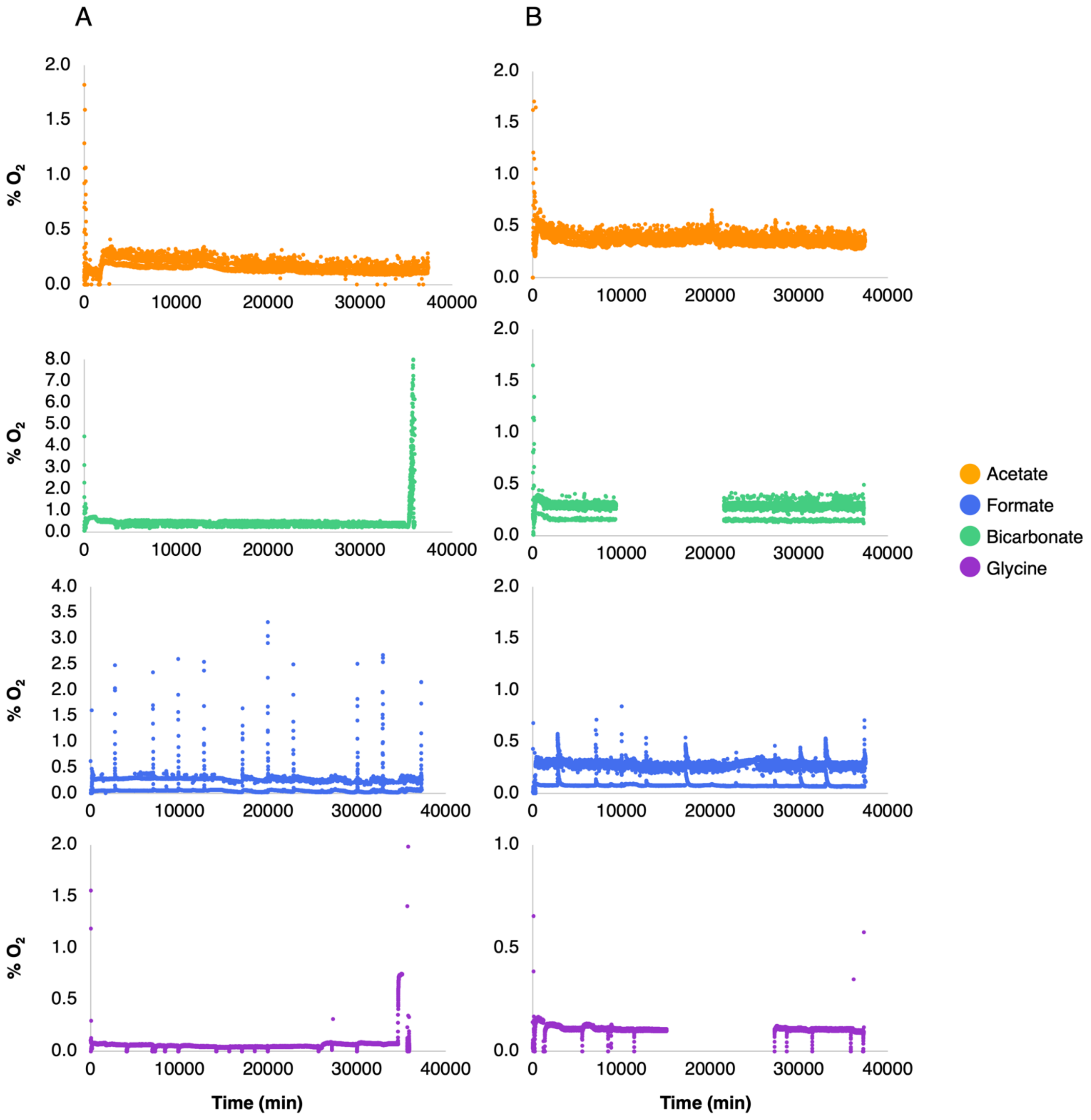


**Supplementary Figure 4.** Percentage of oxygen in the bioreactor gas phase during (A) the first and (B) the second experimental run. In the second replicate of the culture grown on bicarbonate and glycine, data acquisition was temporally interrupted due to a technical problem. In the cultures grown on formate, the oxygen peaks correspond to sampling timepoints. Initial and final oxygen peaks in all cultures correspond to the startup and shutdown of the bioreactors and do not influence the samples discussed in the article.

**Supplementary Table 1.** Average alpha diversity and evenness values per culture condition (see Figure 2). Indices are calculated to the observed number of ASVs, the Shannon and Pielou index in bacterial (A) and archaeal (B) sequences from the inoculum (n=2) and the microbial consortia grown overtime on acetate (n=6), formate (n=6), bicarbonate (n=6) or glycine (n=5). Values obtained from both experimental runs are combined for each culture condition (average ± standard deviation).

| | **A** | | | **B** | | |
|---|---|---|---|---|---|---|
| | **Observed** | **Shannon** | **Pielou** | **Observed** | **Shannon** | **Pielou** |
| **Inoculum** | 315 ± 56.75 | 0.95 ± 0.01 | 4.17 ± 0.34 | 10 ± 4.42 | 0.76 ± 0.04 | 1.75 ± 0.28 |
| **Glycine** | 215.8 ± 42.66 | 0.8 ± 0.11 | 2.93 ± 0.51 | 12.4 ± 7.54 | 0.79 ± 0.03 | 1.74 ± 0.25 |
| **Bicarbonate** | 76.17 ± 69.85 | 0.69 ± 0.22 | 2.07 ± 1.11 | 8.4 ± 4.51 | 0.72 ± 0.09 | 1.5 ± 0.3 |
| **Formate** | 80.5 ± 78.8 | 0.61± 0.32 | 1.86 ± 1.19 | 8.2 ± 2.86 | 0.75 ± 0.12 | 1.63 ± 0.38 |
| **Acetate** | 72.5 ± 95.65 | 0.47 ± 0.39 | 1.54 ± 1.59 | 6.6 ± 1.52 | 0.75 ± 0.02 | 1.52 ± 0.1 |
